# Analysis of the 4q35 chromatin organization reveals distinct long-range interactions in patients affected with Facio-Scapulo-Humeral Dystrophy

**DOI:** 10.1101/462325

**Authors:** Marie-Cécile Gaillard, Natacha Broucqsault, Julia Morere, Camille Laberthonnière, Camille Dion, Cherif Badja, Stéphane Roche, Karine Nguyen, Frédérique Magdinier, Jérôme D. Robin

## Abstract

Facio-Scapulo Humeral dystrophy (FSHD) is the third most common myopathy, affecting 1 amongst 10,000 individuals (FSHD1, OMIM #158900). This autosomal dominant pathology is associated in 95% of cases with genetic and epigenetic alterations in the subtelomeric region at the extremity of the long arm of chromosome 4 (q arm). A large proportion of the remaining 5% of cases carry a mutation in the *SMCHD1* gene (FSHD2, OMIM #158901). Here, we explored the 3D organization of the 4q35 locus by three-dimensions DNA *in situ* fluorescent hybridization (3D-FISH) in primary cells isolated from patients. We found that D4Z4 contractions and/or *SMCHD1* mutations impact the spatial organization of the 4q35 region and trigger changes in the expression of different genes. Changes in gene expression was corroborated in muscle biopsies suggesting that long distance interaction modulate expression of a number of genes at the 4q35 locus in FSHD. Using induced pluripotent stem cells (hIPSC), we examined if chromatin organization is inherited or acquired during differentiation. Our data suggest that folding of the 4q35 region is modified upon differentiation. These results together with previous findings highlight the role of the D4Z4 repeat in the organization of chromatin and further indicate that the D4Z4-dependent 3D structure induces transcriptional changes of 4q35 genes expression.

## Introduction

Over the recent years, sequence distribution within the nuclear space, nuclear topology and long distance interactions have emerged as key elements in the regulation of gene expression and cell fate determination. The genome is separated from the cytoplasm by the nuclear envelope. The inner nuclear membrane is composed of specialized proteins, which contribute to the higher-order organization of the genome within the nuclear space and the regulation of gene expression, with repressed loci mainly located at the nuclear periphery (Andrey and Mundlos, 2017; Robin and Magdinier, 2016).

Interphase chromosomes reside in minimally overlapping chromosome territories with active genes mainly oriented toward the nuclear interior (T. Cremer and C. Cremer, 2001). Individual genes are largely confined to their respective chromosome territories (CT). However, in certain developmental contexts, certain loci such as the *Hox* genes or genes on the inactive X chromosome can loop out of their CTs (Chambeyron and Bickmore, 2004; Chaumeil et al., 2006). Furthermore, long distance dynamic interactions between sequences or subnuclear domains have been identified by various genome-wide approaches (Jerković et al., 2017; Stavreva et al., 2015; Y. Wang et al., 2018). Thus, different topological levels shape the human genome, from large scale folding involving *cis-* or *trans*-long-distance interactions to smaller interactions between regulatory elements located in close proximity (Robin and Magdinier, 2017). Recent data suggest a high level of topological conservation between cell types and species (Lieberman-Aiden et al., 2009; Rao et al., 2014). However, only a few sequences with topological activity have been identified and characterized (Ottaviani et al., 2010; 2009; Zullo et al., 2012). Amongst them, repeat elements linked to pathologies have been recently associated with topological modifications (Sun et al., 2018).

Regarding this topological organization, one intriguing locus is the 4q35 subtelomeric region (Masny et al., 2004; Tam et al., 2004). This gene-poor locus characterized by the presence of gene-deserts and large blocks of repetitive DNA sequence (e.g., a 3.3Kb macrosatellite unit dubbed D4Z4, among others) is localized at the nuclear periphery, a nuclear compartment enriched in heterochromatin (Masny et al., 2004; Ottaviani et al., 2009; Reik, 2007; Tam et al., 2004; Ueda et al., 2014). 4q35 genes are organized in clusters separated by domains associated with the nuclear lamina (Lamin Attachment Domains, LAD) (Gonzalez-Suarez et al., 2009; Guelen et al., 2008), limited by CTCF-dependent boundaries. We have previously described the D4Z4 macrosatellite on the 4q35 region as the first element able to tether any telomere at the nuclear periphery and control the replication timing of its abutting telomeric region. In general, domains located at the nuclear periphery harbor heterochromatin features and correspond to silenced regions (Guelen et al., 2008; Ikegami et al., 2010). As usually observed for heterochromatin-rich regions, the 4q35 telomere replicates late, at the end of the S phase (Arnoult et al., 2010) but displays features of repressed euchromatin rather than constitutive heterochromatin (Tsumagari et al., 2008).

Interestingly, shortening of the D4Z4 array on the 4q35 locus is linked to Facio-Scapulo-Humeral Dystrophy (FSHD), the third most common hereditary myopathy. In most cases (95%) FSHD is linked to the deletion of an integral number of D4Z4, on one of the two 4q35 alleles (Sarfarazi et al., 1992; Wijmenga et al., 1993). Indeed, affected individuals exhibit between 1-10 repeats whereas the general population have a number of repeats above 10 (up to 100) (Lemmers et al., 2010). Shortening of the array is associated with DNA hypomethylation and relaxation of the macrosatellite chromatin suggesting the involvement of epigenetic changes in the disease (Gaillard et al., 2014; Jones et al., 2015; Robin et al., 2015; Stadler et al., 2013). In agreement, the remaining 5% of patients usually display a profound hypomethylation and 80% of them carry a mutation in gene encoding the SMCHD1 (Structural Maintenance of Chromosomal Hinge Domain Containing 1) protein (Lemmers et al., 2012). In mice, *Smchd1* loss of function results in early lethality for female embryos, attributed to derepression of genes on the inactive X chromosome (Blewitt et al., 2008). Smchd1 is also involved in silencing of repetitive DNA sequences and formation of long distance loops at the *Hox* genes loci and inactive X chromosome (Jansz et al., 2018).

By using a three-dimensional Fluorescent *in Situ* hybridization approach (3D FISH), we have previously shown the existence of functional interactions between D4Z4, the nuclear lamina and the telomere (Arnoult et al., 2010; Ottaviani et al., 2010; 2009; Robin et al., 2015) with a key role for D4Z4 in the organization and regulation of long-distance interactions at this locus. These observations were further corroborated by others, with chromatin conformation capture approaches that revealed the influence of the contracted units on the 3D organization with chromatin loops encompassing domains beyond the D4Z4 units (Bodega et al., 2009; Petrov et al., 2006; Pirozhkova et al., 2008). The higher-order regulation of this region is also modulated by telomere length (Robin et al., 2015), suggesting possible cooperation between different specialized genomic elements in the organization of this region. Hence, partitioning of the 4q35 region into different topological domains raises the question of the regulation of long distance interactions across the locus.

In order to get a precise insight into the 3D DNA interactions within the 4q35 locus in normal and pathological conditions, we investigated the three-dimensional chromatin organization of the locus using hybridization techniques in primary cells isolated from FSHD individuals. We identified different loops organizing the 4q35 locus into different chromatin domains. In agreement with previous findings and a role for D4Z4 in chromatin topology, these chromatin domains are modified when the number of D4Z4 units is decreased. Moreover, we analyzed expression of a number of genes localized within or outside of these loops in our cells and confronted our results to those obtained in muscle biopsies from affected and unaffected individuals.

## Results

### Exploring the 3D organization of the 4q35 locus

Given the peculiar organization of the 4q35 region and the topological role played by D4Z4 in healthy and diseased cells, we aimed at deciphering the topological changes of this locus in FSHD1 and 2 patients compared to healthy donors. To this aim, we first analyzed datasets available in the literature (Y. Wang et al., 2018). Indeed, In the past decade, a wealth of chromatin techniques coupled to deep-sequencing technologies have emerged and multiple databases on the 3D DNA organization have been created (Dixon et al., 2012; Rao et al., 2014; Schmitt et al., 2016). We retrieved 4q35 locus interaction maps (183-191Mb of Chr. 4) generated from 3 different cell lines and 1 biopsy (Supplementary Figure 1). Amongst these maps, two achieved a Kb resolution (GM12878, IMR90) while the remaining two barely achieved a Mb resolution (STL002, *Psoas* muscle biopsy), a poor resolution mostly due to the depth of sequencing, number of assay performed and nature of the samples.

Altogether these maps show a conserved domain organization of the most subtelomeric part of the locus up to a region located between the *FAT1* and *SORBS2* genes (187 and 186 Mb of Chr.4; respectively), two genes previously implicated in FSHD (Caruso et al., 2013; Puppo et al., 2015; Robin et al., 2015).

Given the unique genomic rearrangement of this subtelomeric region, we next set out to investigate the chromatin organization of the 4q35 locus by 3D DNA FISH using selected probes in well-defined cells (Table 1). We used primary fibroblasts isolated from skin biopsies from two different FSHD1 patients, one of which presenting a homozygous D4Z4 contraction (i.e., 8 UR) along with fibroblasts isolated from two FSHD2 individuals (*i.e.*, >10 UR) carrier of a mutation in *SMCHD1* (Table 1).

**Table 1.** Summary of cells (fibroblasts from skin biopsies) used in the study with their respective associated status and number of D4Z4 repeats (RU) at the 4 and 10q locus along with mutation found in SMCHD1. FSHD individuals were diagnosed by a clinical examination by neurologists with clinical expertise in the evaluation of neuromuscular diseases. The genetic status was confirmed by genetic testing (Southern Blot, DNA combing). Gender and age at biopsy are indicated as well as the corresponding identification number (ID) of samples used elsewhere (Dion et al., under Review)

| Sample | Status | 4q | 10q | Age | Sex | hIPSCs | SMCHD1 |
| --- | --- | --- | --- | --- | --- | --- | --- |
| A | Control | >10RU | NA | Foetus | ♀ | Available | WT |
| N<br>(ID: 12759) | FSHD1 | 7RU qA<br>>10RU qB | NA | 51 | ♀ | Available | WT |
| H | FSHD1 | 8RU qA<br>8RU qA | NA | 29 | ♂ |  | WT |
| G<br>(ID: 11440) | FSHD2 | 12RU qA<br>32RU qA | 14UR qA<br>17UR qA | 37 | ♂ |  | c.2338+4A>G intron<br>18 r.2261_2337del;<br>p.S754* |
| S<br>(ID: 15166) | FSHD2 | 12RU qA<br>23RU qB | 13UR qB<br>20UR qB | 67 | ♀ |  | c.4614_4615<br>insTATAATA;<br>r.4614_4615<br>insTATAATA;<br>p.A1539Yfs*4 |

As no clear TADs were found in more centromeric regions and given the putative interactions between the FSHD locus and the nuclear lamina or matrix (Masny et al., 2004; Ottaviani et al., 2009; Petrov et al., 2006; Robin et al., 2015; Tam et al., 2004), we focused our attention on the different regions separated by LADs (Guelen et al., 2008). The 4q35 locus contains four LADs. The most distal one (190-191 Mb of Chr 4) is located upstream of the D4Z4 array and overlaps with a large gene-desert between the region containing the *ZFP42* and *FAT1* genes. The second LAD (187-187.7 Mb) separates the *FAT1* gene from the *SORBS2* region and encompasses several genes (including *MTNR1A* and *FAM149A*). The third LAD (184.8-185.4 Mb) is situated in a gene-poor region along with the fourth LAD (185.6-185.8 Mb), suggesting the existence of two loops anchored at the nuclear periphery between the *ACSL1* and *WWC2*-containing domains (Kent et al., 2002). Interestingly, the *WWC2* region correspond to a TAD boundary, at least in two data sets (GM12878, IMR90; respectively)(Y. Wang et al., 2018). To understand higher-order 3D folding patterns of the 4q35 region in the different contexts, we selected five probes encompassing the whole 4q35 regions and the different putative topological domains.

### 3D genome organization of the 4q35 locus is modified in FSHD

Single cell FISH analysis of genes or entire chromosomes shows that individual cell exhibit the same genomic organization regardless of cell types (Meaburn et al., 2009; Rao et al., 2015). To investigate the three-dimensional organization of the 4q35 region of each individual allele in FSHD, we thus performed 3D DNA FISH in FSHD1 or 2 fibroblasts and control, using for each assay two sets of probes (Figures 1-2, Supplementary Figures 2-5). One probe was common between all experiments and hybridize with the D4S139 minisatellite (red) proximal to the D4Z4 repetitive array and corresponding to the most proximal region specific to the 4q35 locus. Rationale for the choice of remaining probes is described above and targeted the *FAT1, SORBS2, ASCL1* and *WWC2* regions, respectively (green). After hybridization and confocal imaging, stacks of nuclei were processed using the IMARIS software (Bitplane, AG) for 3D reconstruction and analysis (Supplementary Figure 2). Three Dimensions computational modeling allows one to retrieve accurate values from confocal imaging such as volume, distribution, position and distances. We compiled results for each assay and asked whether 4q35 folding is identical between cells from controls and patients and between FSHD1 and FSHD2.

**Figure 1.**
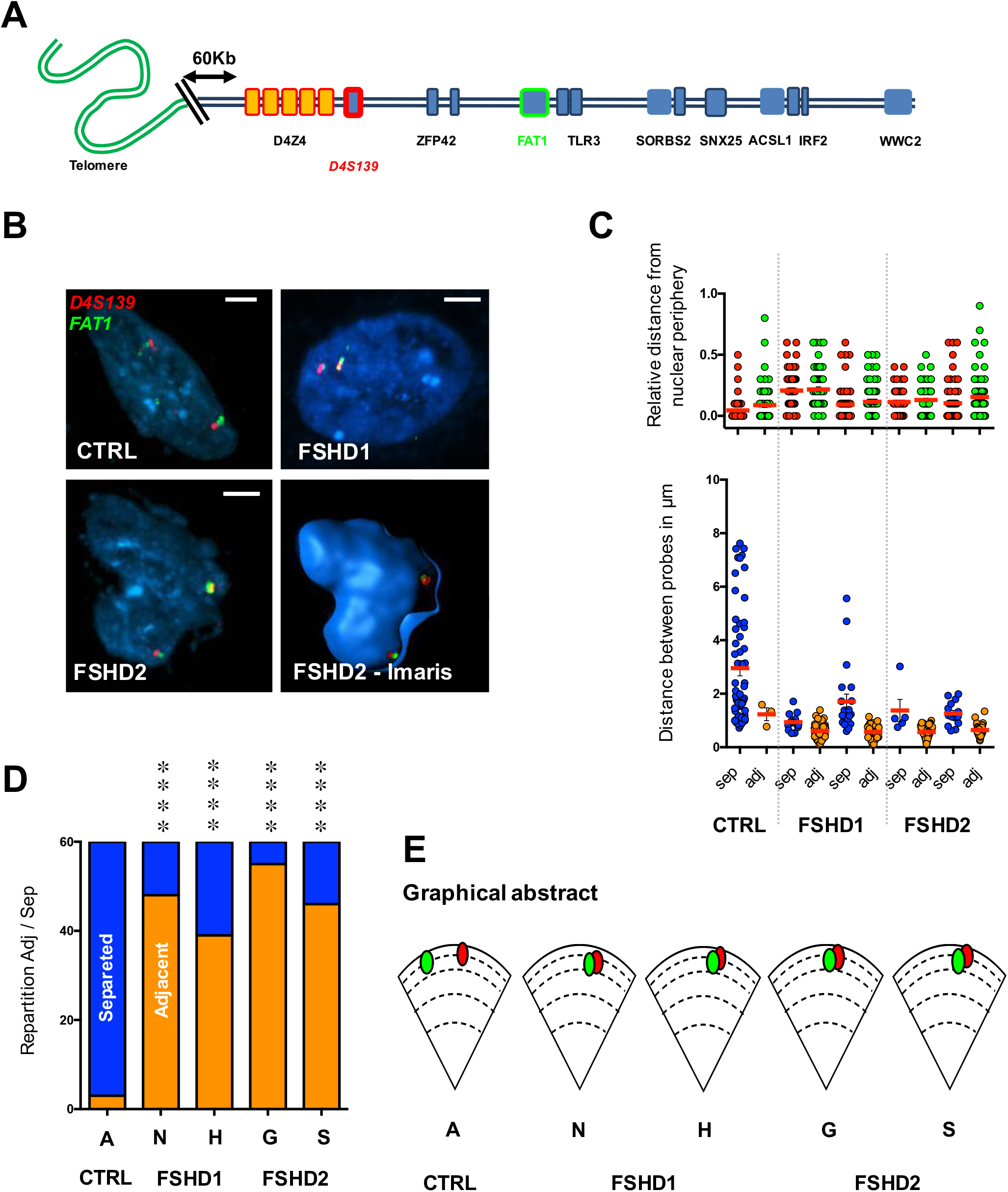
The distance between the distal 4q35 region and the *FAT1* gene is decreased in cells from FSHD1 and 2 patients. **A.** Graphical representation of the subtelomeric region of chromosome 4, from the telomere (green) to the *WWC2* gene (blue square) located at a distance of 7 Mb from the D4Z4 array. 3D FISH was done using two sets of probes corresponding either to the D4S139 variable number tandem minisatellite (VNTR; red) or the *FAT1* region (green). **B**. Representative pictures of FISH images in primary fibroblasts from healthy donors (control) or FSHD1 and FSHD2 patients and 3D reconstruction using IMARIS. Scale is indicated by a white bar (=2µm) **C.** Quantification of FISH signal was done using IMARIS. We measured the distance of each probe to the nuclear periphery (upper graph) and distance between the probes (lower graph). At least 30 nuclei were analyzed (60 alleles). For each cell, we evaluated if signals were separated or adjacent (sep, adj; respectively). **D.** Summary of events and associated Chi-Square test (n=30 cells per sample; 60 alleles). Frequency of adjacent signals is increased in all FSHD cells. **** *p* < 0.001. **E.** Summary of the average distances between probes and their respective distances to the periphery in each sample. Signals are mostly localized at the periphery regardless of disease status.

First, all signals were found at the nuclear periphery regardless of disease status and D4Z4 contractions (average distance from nuclear periphery <0.3µm; Figures 1-2). Next, we evaluated if signal between probes were separated or adjacent (sep, adj; respectively). Our binary decision (sep, adj) was based on the distances measured between the gravity center of the probe signal after 3D reconstruction (Supplementary Figure. 2). Overall, distances associated to adjacent and separated events were significantly different between the conditions tested (p<0.0001; Mann Whitney, *α*=0.05); signals were considered as adjacent when the measured distance was below 0.9µm (Mean ± SD; adj distance = 0.60 ± 0.31µm). In control cells, the D4S139 region is localized in the vicinity of the *SORBS2* and *WWC2* regions, but separated from *FAT1* and *ASCL1* (all in different LADs), a finding reminiscent of the TAD identified in the literature (Supplementary Figure 1)(Y. Wang et al., 2018). In FSHD cells, Interactions were identical between FSHD1 and 2 in all conditions but one (D4S139-*SORBS2*; Figure 2A). The D4S139 region no longer interacts with *WWC2* but with *FAT1* (p<0.001 for FSHD cases; Chi-Square test, *α*=0.05), suggesting that the topological organization of this locus is orchestrated by epigenetic features associated with the D4Z4 array and its peripheral subnuclear localization.

Strikingly, regarding *SORBS2*, we observed a partial loss of interaction restricted to FSHD1 cells. Indeed, when considering FSHD1 cells carrying only one contracted allele (Table 1, N), 85% of cells displayed a partial loss of interaction where only one out of two signals were separated (Figure 2A). Accordingly, in FSDH1 cells carrying two contracted alleles (Table 1, H), the interaction was completely lost (Figure 2, p <0.001; Chi-Square test, *α*=0.05) suggesting that this interaction is alleviated when the number of D4Z4 repeats is decreased as previously observed (Robin et al., 2015).

**Figure 2.**
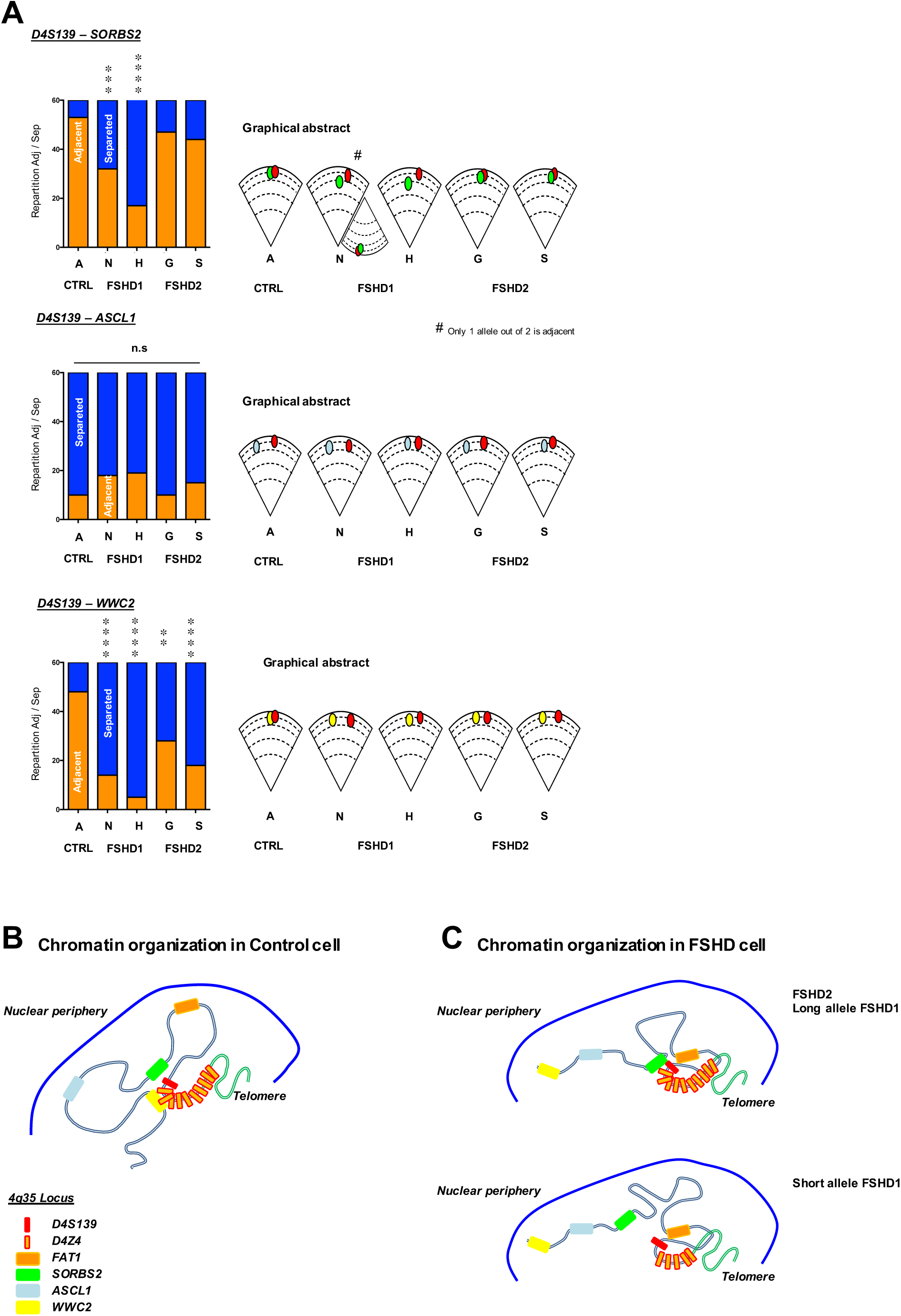
Distinct long distance interactions at the 4q35 region in controls and FSHD cells. **A.** Quantifications of colocalization and nuclear distribution of 3D DNA FISH assay within the 4q35 locus. We report the frequency of association between the D4S139 region (red; as in Figure 1) and either the *SORBS2, ASCL1* or *WWC2* region (green, blue, yellow; respectively) in primary fibroblasts from healthy donors, FSHD1 or FSHD2 patients. A Chi-Square test was performed for each assay to determine statistical differences (n=30 cells per sample, 60 alleles). **B-C**. Graphical representation of the 4q35 chromatin organization as revealed by 3D DNA FISH in control cells (**B**) and FSHD1 and 2 cells (**C**). ** *p* < 0.01; *** *p* < 0.005; **** *p* < 0.001.

No difference was found between samples considering the D4S139 and *ASCL1* probes, for which signals were separated in all situations. Taken together our results detail the spatial organization of the 4q35 locus within the nucleus and reveal the existence of new topological domains, dependent of the number of D4Z4 repeats (Figure. 2B-C).

### Changes in chromatin organization of the 4q35 locus correlate with modulation in gene expression

To further investigate the consequences triggered by changes in DNA folding within the 4q35 region, we examined the level of expression of different genes located within or outside the chromatin loops by RT-qPCR (Figure 3). In brief, we focused our attention on genes localized in loops present in control cells but absent in FSHD cells and *vice versa*. Interestingly, expression of most genes within the last 7Mb of chromosome 4 was significantly modulated, with a high variability between samples (e.g., excluding *FAT1* and *STOX2*, p>0.05; Kruskal-Wallis multiple comparison test; *α*=0.05).

**Figure 3.**
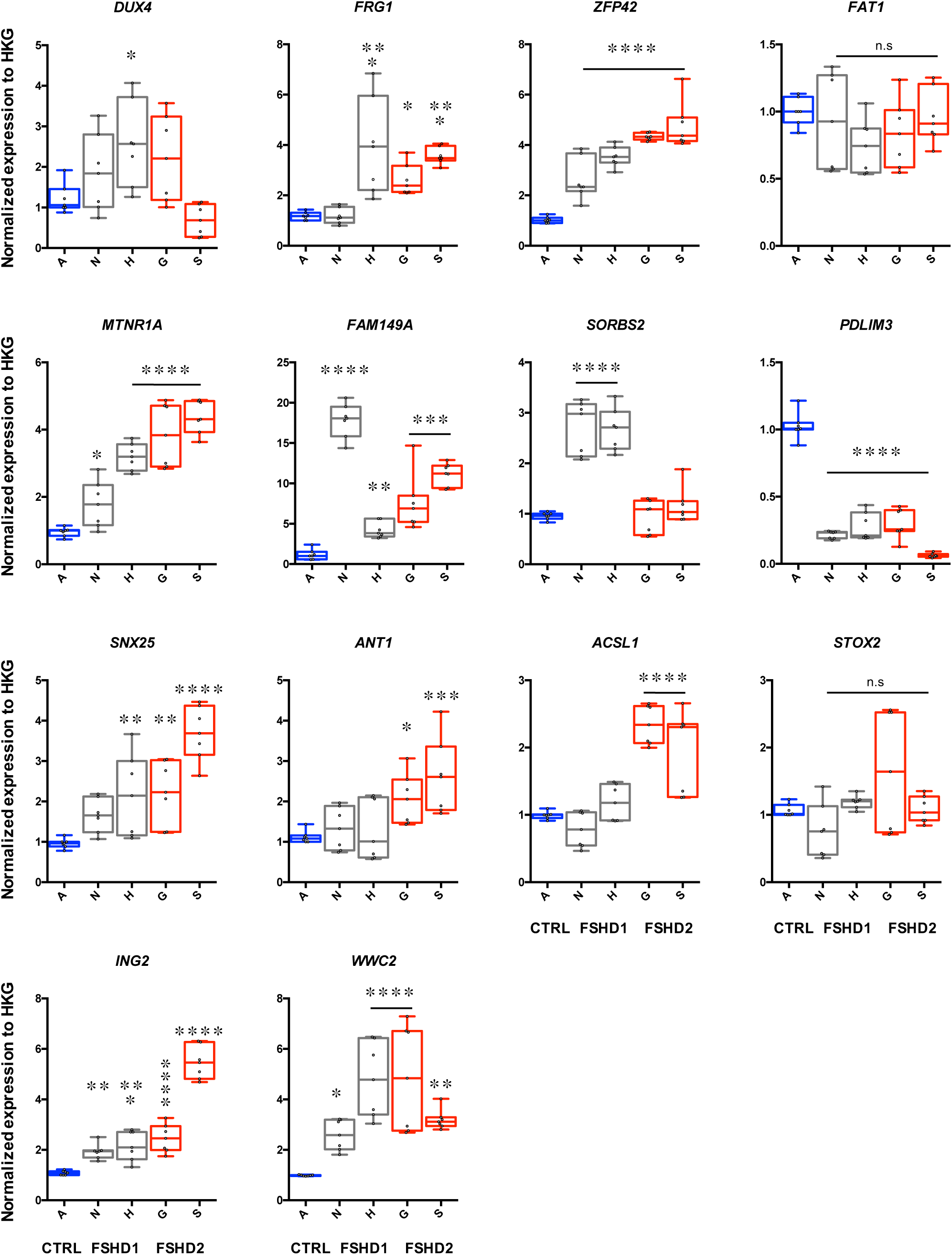
Expression of genes of the 4q35 region in controls and FSHD cells. Gene expression quantified by RT-qPCR in controls, FSHD1 (N, H) and FSHD2 (G, S) cells. Each measure represents the average fold-change expression of six independent assays (biological triplicate in technical RT duplicate) normalized to three housekeeping genes (HKG: *HPRT, PPIA* and *GAPDH*; △△Ct method). Differences compared to the control cells are represented as boxplot with associated statistical significance (Kruskal-Wallis multiple comparison test; *α*=0.05) along with means ± SEM. * *p* <0.05; ** *p* < 0.01; *** *p* < 0.005; **** *p* < 0.001.

First, regarding the D4S139-*FAT1* loop (Figure 2C); we detected an upregulation of the *FRG1* and *ZFP42* genes in all FSHD cells regardless of the D4Z4 status (FSHD1 and FSHD2) (p<0.05; Kruskal-Wallis multiple comparison test; *α*=0.05). However, one FSHD1 sample was not significantly different from the control cells regarding *FRG1* expression (p>0.9; Kruskal-Wallis multiple comparison test; *α*=0.05).

Next, we deduced an interaction between *SORBS2* and *FAT1*, likely restricted to the uncontracted allele of FSHD cells only (N,G,S; Figure 2A). Expression of genes located within this newly formed loop was significantly upregulated not only in FSHD cells presenting the interaction but across all (*MTNR1A, FAM149A*). This suggest other modulators of expression, including a possible unidentified loop (Ghirlando et al., 2012). Strikingly, expression of *SORBS2* was directly correlated to the formation of the loop. Only cells where the D4S139-*SORBS2* interaction was not detected presented upregulation of its expression (FSHD1; N, H; p<0.001; Kruskal-Wallis multiple comparison test; *α*=0.05).

Our set of 3D FISH assay did not allow us to detect other chromatin structures beyond the *SORBS2* locus in FSHD cells, whereas control cells showed a D4S139-*WWC2* interaction. Expression of genes beyond the *SORBS2* locus displayed mixed results. The *ING2* and *WWC2* gene were upregulated in FSHD cells (p<0.001; Kruskal-Wallis multiple comparison test; *α*=0.05) while *ANT1* and *ACSL1* were upregulated only in FSHD2 cells.

Collectively, our data link changes in gene expression and modification in chromatin organization. Importantly, if one can not exclude the tissue-specific expression of certain genes, insights from global expression analyses can illustrate modifications in chromatin domains (Dixon et al., 2016; Ghirlando et al., 2012). Hence, we exploited the gene expression analysis to decipher the emergence of additional territories (Supplementary Figure 6). We used the ratio between housekeeping genes and the gene of interest for normalization. We found that in FSHD cells, levels of expression were globally increased beyond the *SORBS2* locus, advocating for a looser chromatin structure allowing the binding of the transcription machinery. Last, we did not detect *DUX4* overexpression in FSHD cells (H, p=0.0176; Kruskal-Wallis multiple comparison test; *α*=0.05) likely due to the type of cells (isolated primary fibroblast cells).

### 4q35 genes are differentially expressed in muscle biopsies

In primary fibroblasts, we found that the modification of the 4q35 chromatin landscape correlates with changes in transcription of genes found within the modified domains. As a second step, we thus consolidated our findings by analyzing gene expression directly in muscle biopsies isolated from individuals diagnosed with either FSHD1 or FSHD2. Hence, for confronting our *in vitro* data with tissues affected in this pathology (i.e., skeletal muscle), we used a collection of biopsies isolated from control and affected individuals (FSHD1, n= 7; FSHD2, n= 3; Control, n= 7; Table 2). As found by others, *DUX4* and *FRG1* expression were upregulated in FSHD biopsies (p<0.005; Kruskal-Wallis multiple comparison test; *α*=0.05) (Broucqsault et al., 2013; Gabellini et al., 2002; Lemmers et al., 2010). Additionally, *FAM149A* and *SORBS2* were exclusively upregulated in FSHD1 (p<0.001, respectively; Kruskal-Wallis multiple comparison test; *α*=0.05); *FAT1* and *ASCL1* in FSHD2 (p=0.023 and p=0.006, respectively; Kruskal-Wallis multiple comparison test; *α*=0.05). If one considered that chromatin territories are conserved throughout cell types, our results strongly suggest a direct correlation between 3D organization of the 4q35 locus (e.g., no interaction in FSHD1) and *SORBS2* or *WWC2* expression in FSHD. Moreover, the intriguing upregulated expression of *ASCL1* and *FAT1* restricted to FSHD2 biopsies advocates for a more complex regulation possibly involving SMCHD1 as described for other genomic regions (Jansz et al., 2018; C.-Y. Wang et al., 2018), or the existence additional DNA interaction yet to be characterized (Robin et al., 2015).

**Table 2.** Summary of muscle biopsies used in the study with their respective status. FSHD individuals were first diagnosed by a clinical examination (except for fetuses obtained after medical abortion) and further confirmed by genetic testing (Southern Blot, DNA combing). Muscle origin, gender and age at biopsy are reported.

| Sample | Status | Origin | Age | Sex |
| --- | --- | --- | --- | --- |
| 1 | Control | Quad | 77 | ♂ |
| 2 | FSHD1 | Quad | Fetus | ♀ |
| 3 | Control | Quad | Fetus | ♂ |
| 4 | Control | Quad | Fetus | ♂ |
| 5 | FSHD1 | Quad | 61 | ♂ |
| 6 | FSHD2 | Quad | 57 | ♂ |
| 7 | FSHD2 | Quad | 45 | ♂ |
| 8 | FSHD2 | Quad | 42 | ♂ |
| 9 | FSHD1 | Quad | 54 | ♂ |
| 10 | Control | Quad | 52 | ♂ |
| 11 | Control | Quad | Fetus | ♂ |
| 12 | FSHD1 | Quad | 61 | ♂ |
| 13 | Control | Quad | 69 | ♂ |
| 14 | Control | Quad | 71 | ♀ |
| 15 | FSHD1 | Quad | Fetus | ♂ |
| 16 | FSHD1 | Quad | Fetus | ♀ |
| 17 | FSHD1 | Quad | Fetus | ♀ |

### The 4q35 chromatin organization is set to a new conformation in hIPSC

Earlier works have shown that the position of single gene loci and entire chromosomes is heritable after cell division (Strickfaden et al., 2010). TADs are largely maintained through development as TAD boundaries tend to be 65% similar among cell types (Phillips-Cremins et al., 2013; Rao et al., 2015; Vietri Rudan et al., 2015). However, HiC data revealed massive transformation in chromosome topology after reprogramming with many interactions present in the cells of origin being progressively erased from early to late passage (Gonzales and Ng, 2016). Furthermore, the nuclear lamina undergoes profound changes since A-type Lamins present in the somatic cell-of-origin are not expressed in pluripotent cells (Constantinescu et al., 2006) suggesting possible topological changes in the 3D organization of the 4q35 locus in pluripotent cells. We therefore investigated whether folding was inherited or modified after reprogramming. Taking advantage of the chromatin organization displayed in FSHD1 cells carrying both contracted and uncontracted *D4Z4* allele (N, Table 1), we restrained our 3D DNA FISH analysis to hIPSCs generated from the corresponding fibroblasts (A and N; respectively). HIPSCs cells are described elsewhere (Badja et al., 2014) (Dion et al., under review). We analyzed the 4q35 conformation using probes for the D4S139 region together with probes targeting the *FAT1, SORBS2, ACSL1* and *WWC2* loci (Figure 4). Remarkably, no differences were found between control and FSHD1 hIPSCs. Signals were all localized at the nuclear periphery and only one long-distance interaction was preserved after reprogramming between D4S139 and *SORBS2* (Figure 4B), consistent with the main TAD found in other cell types. In parallel, we tested gene expression by RT-qPCRs (Figure 5). Our data revealed no significant differences between conditions, expect for the *DUX4* gene upregulated in FSHD1 hiPSCs (p=0.0130; Mann Whitney test; *α*=0.05). Thus, our data suggest that upon reprogramming, cells undergo drastic chromatin structure modification within the 4q35 locus. This pluripotent state display a domain encompassing the most subtelomeric part of the locus and the *SORBS2* region which appears to be the most conserved domain among cell types as also revealed by HiC (Supplementary figure 1). To note, as seen by others, *DUX4* expression was independent of the centromeric 4q35 chromatin organization, but might be under the influence of other chromatin features such as changes in DNA methylation, attachment to the nuclear matrix or telomere length (Gaillard et al., 2014; Petrov et al., 2006; Stadler et al., 2013).

**Figure 4.**
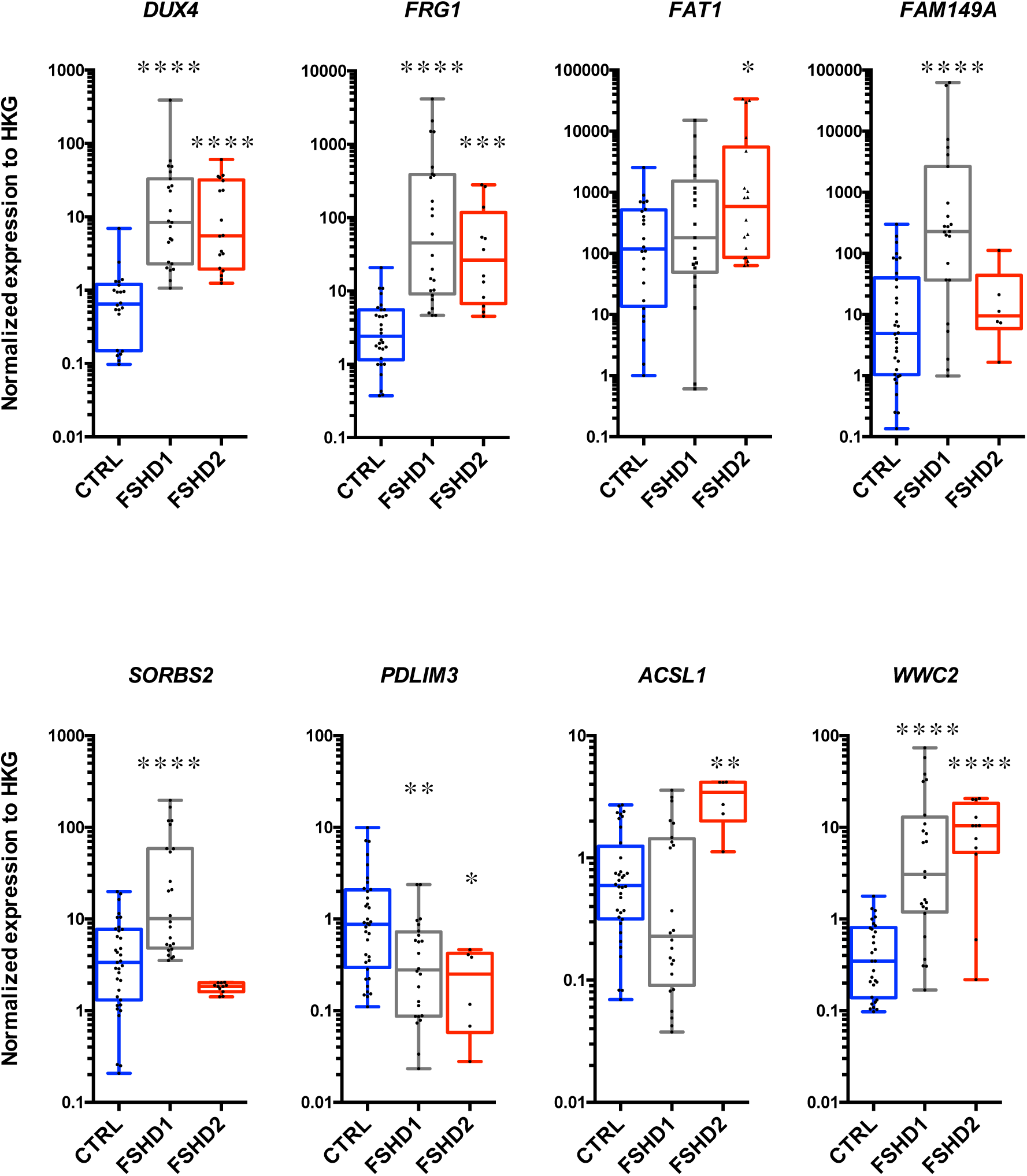
Expression of genes of the 4q35 region in controls and FSHD biopsies. Gene expression quantified by RT-qPCR in control (n=7, Blue); FSHD1 (n=7, Grey) and FSHD2 (n=3, Red) muscle biopsies (*quadriceps femoris*). Each measure represents the average fold-change expression of six independent assays (RT triplicate, technical duplicate) normalized to three housekeeping genes (HKG: *HPRT, PPIA* and *GAPDH*; △△Ct method). Differences from the control group are represented as boxplots with associated statistical significance (Kruskal-Wallis multiple comparison test; *α*=0.05) along with means ± SEM. * *p* <0.05; ** *p* < 0.01; *** *p* < 0.005; **** *p* < 0.001.

**Figure 5.**
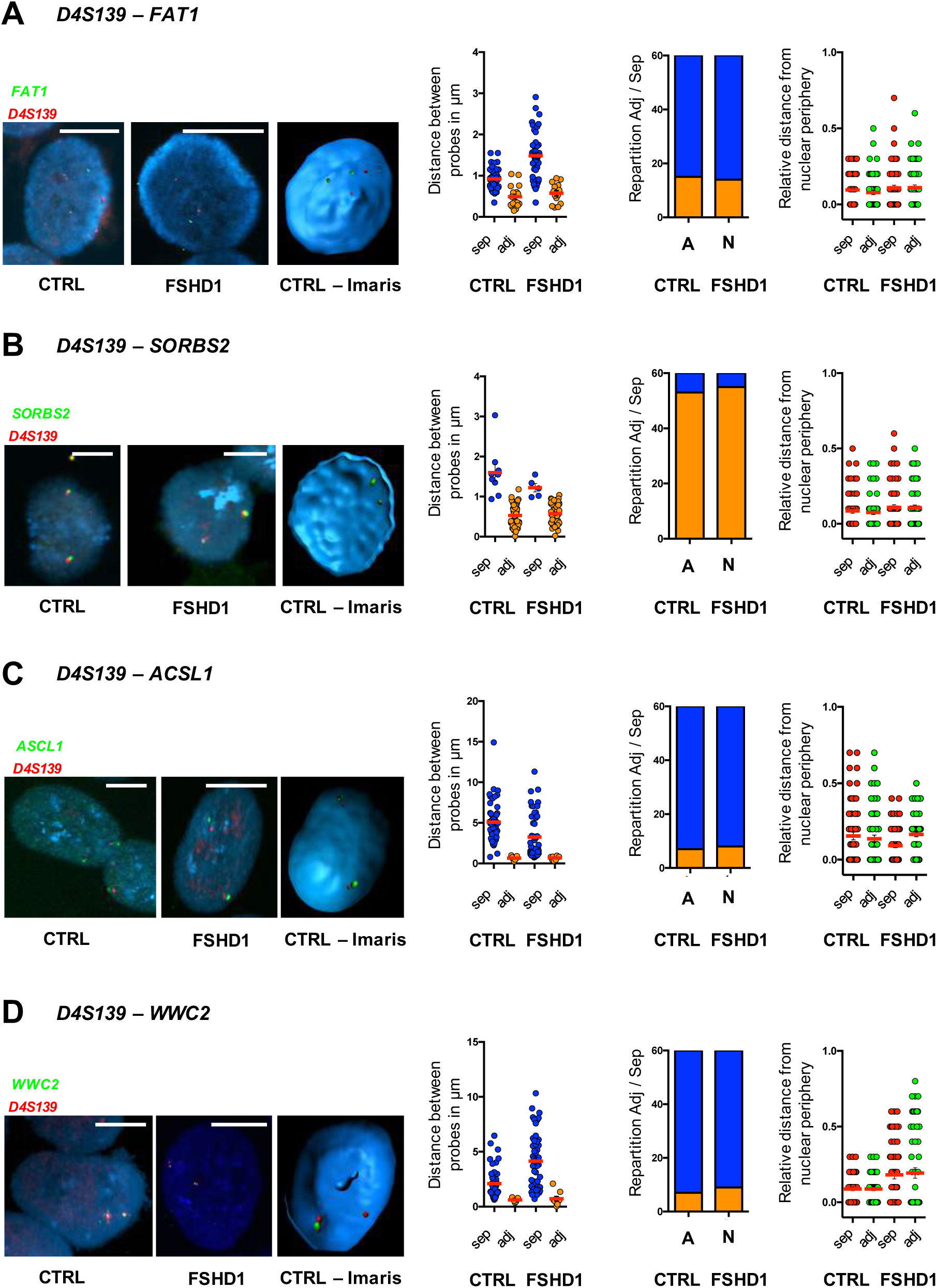
Topology of the 4q35 region is identical in control and FSHD pluripotent stem cells. Three-Dimension DNA FISH assay within the 4q35 locus in human Induced Pluripotent Stem Cells (hIPSC) generated from control and FSHD1 primary fibroblasts (A, N; respectively - see Table. 1). 3D FISH was done using two sets of probes corresponding either to the D4S139 region (red) or different loci along the 4q35 chromosome (green, *FAT1, SORBS2, ASCL1* or *WWC2*; respectively). Representative pictures and 3D reconstruction using IMARIS along with their associated quantifications are presented. We measured the distance of each probe to the nuclear periphery and distance between the probes. For each cell, we evaluated if signals are separated or adjacent (sep, adj; respectively). Scattergram distribution of events and associated Chi-Square test (n=30 cells per sample, 60 alleles) are shown. Scale is indicated by a white bar (=4µm). No statistical differences were found between cells. **A.** Localization between the D4S139 region and the *FAT1* gene. **B.** Localization between the D4S139 region and the *SORBS2* gene. **C.** Localization between the D4S139 region and the *ACSL1* gene. **D.** Localization between the D4S139 region and the *WWC2* gene.

**Figure 6.**
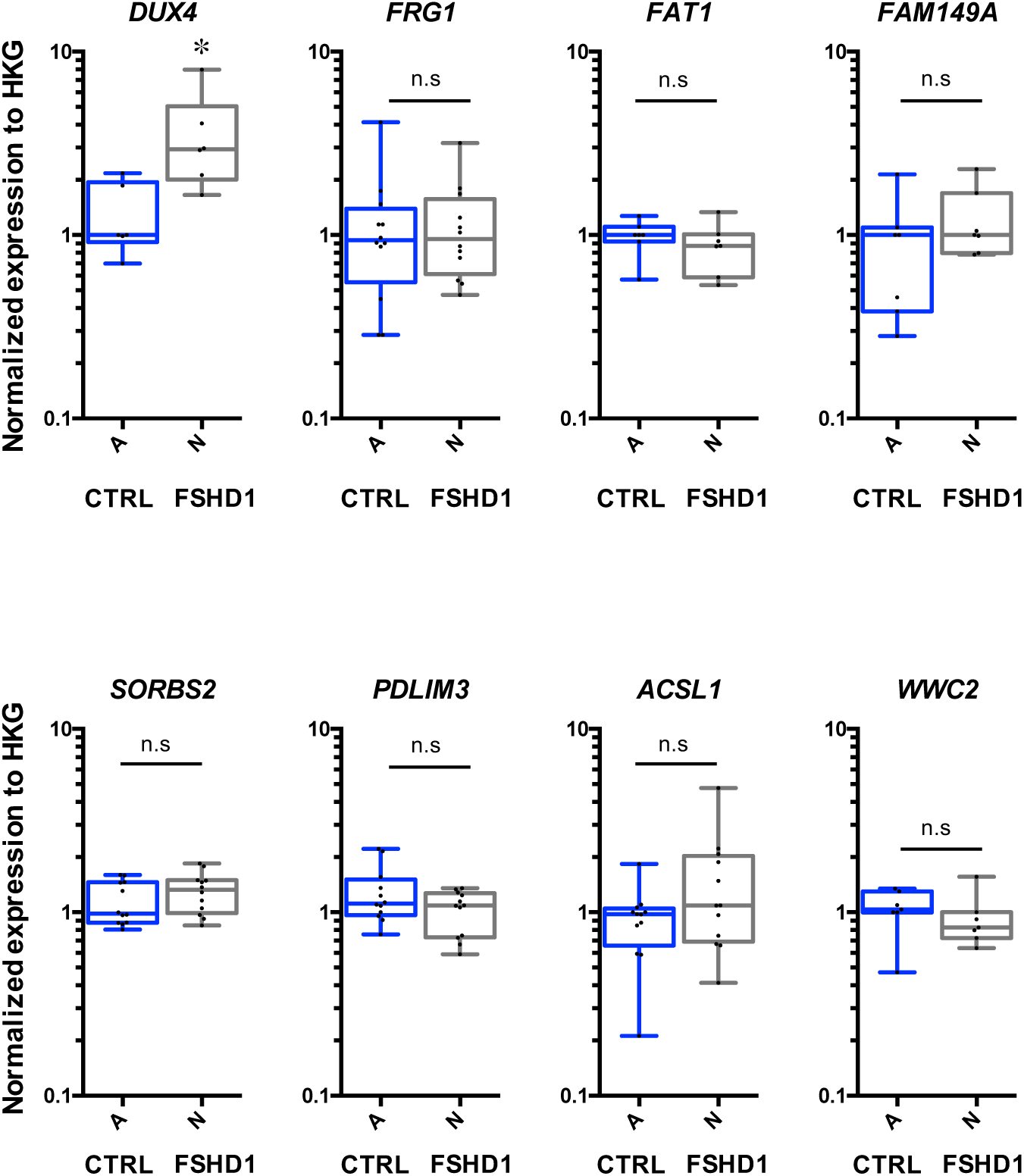
4q35 gene expression is not modulated in FSHD hiPSCs compared to controls. Gene expression quantified by RT-qPCR in control FSHD1 hIPSC (A, N; respectively - see Table. 1). Each measure represents the average fold-change expression of six independent assays (Biological triplicate in technical RT duplicate) normalized to three housekeeping genes (HKG: *HPRT, PPIA* and *GAPDH*; △△Ct method). Differences relative to control cells are represented as boxplots with associated statistical significance (Mann Whitney; *α*=0.05) along with Means ± SEM. * *p* <0.05.

## Discussion

In this study, we investigated the organization of the 4q35 locus linked to Facio-Scapulo Humeral Dystrophy in different contexts and different cell types. Overall, our data shows important 3D changes within the 4q35 locus due but not limited to D4Z4 array contraction. We observed differences in chromatin organization between cells from controls and individuals with clinical signs of the disease (e.g., FSHD1, FSHD2). While one interaction was dependent of the D4Z4 array (e.g., D4S139-*SORBS2*), most interactions characterized are independent of the number of D4Z4 repeats (FSHD2 cases, Figures 1-2). This chromatin landscape is profoundly modified upon reprogramming with erasure of the parent-of-origin architectural organization and establishment of a novel pluripotent-specific profile identical between controls and FSHD1 cells (Figure 5). In somatic cells, modified long-distance interactions correlates with different expression signatures. This observation was validated in muscle biopsies (Figure 4) and advocates for a conserved topology among cell types and a role for D4Z4 or SMCHD1 in this higher-order organization, with consequences on gene expression. We also observed that expression of *DUX4* was independent of changes occurring in the chromatin organization of more centromeric territories, hence confirming findings by others (Bodega et al., 2009; Gaillard et al., 2014; Jones et al., 2015; Pirozhkova et al., 2008; Stadler et al., 2013).

Importantly, our work based on confocal imaging allows one to analyze a per cell and per allele organization of the chromatin, revealing precise frequency and representation of events across multiple cell samples and for each individual allele within a cell. This technical design limits overrepresentation of discrete events that might bias data interpretation as sometimes observed using 3C techniques (Pirozhkova et al., 2008). Likewise, previous work based on DNA replication timing experiments revealed high similarities between samples (97.3%) and cell types (>80%) without any difference for the 4q35 locus between samples (Pope et al., 2011). Those conflicting observations (interactions vs. no interactions) could arise from difficulties inherent to the analysis of subtelomeric regions in general (i.e., GC-rich, late replication), insufficient depth of sequencing and incomplete assembly of subtelomeric sequences (Fudenberg and Imakaev, 2017; Giorgetti and Heard, 2016; Sims et al., 2014). Altogether, these works using sequencing techniques highlight the challenge and raise new questions concerning the peculiar organization of the 4q35 loci in FSHD, where one needs to consider the specificity of each allele. By using only samples from well characterized cells with regard to number of repeats (analyzed by Southern blot or DNA combing)(Nguyen et al., 2017), these concerns were carefully considered in the design of our study.

Boundary elements or insulators are critical for the establishment or maintenance of the genome architecture. Boundary elements bound by the CTCF protein concur to the formation of TADs and long-distance interactions. At the 4q35 region, we have described D4Z4 as a genomic element acting as a CTCF and A-type Lamins dependent insulator (Ottaviani et al., 2010) and able to tether a telomere at the nuclear periphery (Ottaviani et al., 2009). We have previously identified additional sequences with insulator activity along the 4q35 region, which are not able to tether the 4q35 to the nuclear periphery suggesting a key role for the D4Z4 array in the organization of this region within the nuclear space but also in the establishment of long distance interactions (Morere and Magdinier, unpublished data). In agreement with this hypothesis, we previously showed that in the context of a short D4Z4 array and upon telomere shortening, disruption of a certain chromatin loops modulate the expression but more strikingly splicing of genes such as *SORBS2* or *FAM149A* (Robin et al., 2015). Using different cells, our present work revealed the same interactions (Figure 2, 4) with a loop encompassing the *SORBS2* region and the D4Z4 array, conserved between cells with long D4Z4 array. Interestingly, this loop loosened for one out of the two alleles in FSHD1 fibroblasts is reset in hiPSCs after reprogramming and telomerase reactivation reinforcing the importance of the telomere vicinity in the regulation of this locus (Arnoult et al., 2010; Ottaviani et al., 2009; Robin et al., 2015). Interestingly, this loop does not depend on *SMCHD1* gene dosage, since long distance interactions are maintained in FSHD2 cells carrying a heterozygote truncating mutation. This further argues in favor of a pleiotropic role for this chromatin factor with a role in D4Z4 regulation which likely differs from its topological role in the regulation of *Hox* genes or formation of A and B compartments in X inactivation (Jansz et al., 2018; C.-Y. Wang et al., 2018).

At the 4q35 locus, the territory identified by 3D FISH is reminiscent of a TAD identified by others in different cell types (Y. Wang et al., 2018), advocating for a pivotal role in the setting of boundaries within the 4q35 region. Further, by investigating hIPSCs which do not express A-type Lamins (Constantinescu et al., 2006), we observed that FSHD1 cells did not exhibit the same chromatin organization as founder somatic cells with a loss in the 3D Chromatin structure present before reprograming. This intriguing result reveals that tethering to the nuclear periphery of the 4q35 region is not alleviated by absence of A-type lamins as previously observed using artificially fragmented telomeres (Ottaviani et al., 2009). In this telomerase positive context, the short D4Z4 array (e.g., exhibiting an un-masked CTCF site) harbors a peripheral nuclear position but could be untangled with an abutting long telomere, hence masking the CTCF site and modification of long distance interactions. The total topological re-set towards a new organization common between controls and diseased cells suggests a model where differences in chromatin topology would arise during or after differentiation. Hence, one could hypothesize that the FSHD chromatin landscape is not inherited but progressively installed upon differentiation.

Based on our observations, we propose here a complex orchestration of the 4q35 folding upon differentiation with clear differences between control cells and cells from patients affected with FSHD. Combined with previous findings indicating the reorganization of the 4q35 locus in cells from FSHD patients, defining the dynamic 3D organization of this locus could provide further insights into the understanding of the regulation of this locus and transcriptional signature of 4q35 genes during development and differentiation. Considering the high phenotypical and penetrance variability between patients affected with FSHD, our data propose that differences in the folding of this region might contribute to the variegated expression of a number of genes and variability of symptoms.

## Materials and Methods

### Study samples

All individuals have provided written informed consent for the collection of samples and subsequent analysis for medical research. The study was done in accordance with the Declaration of Helsinki. Controls are randomly selected individuals or patient’s relatives, selected in the same age range and sex representation as patients. Controls are neither carrier of any genetic mutation nor affected by any constitutive pathology. Samples are listed in Tables 1 and 2.

### Reprogramming of human fibroblasts and production of hiPSCs

Human iPSCs were generated as described elsewhere (Badja et al., 2014) (Dion et al. Under review). HiPSCs colonies were picked about 4 to 6 weeks after reprogramming based on their ES cell-like morphology. Colonies were grown and expanded in mTeSR™ 1 medium (StemCells) on dishes coated with Matrigel™ (Corning, cat, No. 354277). The different clones used were fully characterized using classical procedures (Dion et al, under review; (Badja et al., 2014)).

### RNA extraction and quality control

Total RNA was extracted using the RNAeasy kit (Qiagen) following manufacturer’s instructions. Quality, quantification and sizing of total RNA was evaluated using the RNA 6000 Pico assay (Agilent Technologies Ref. 5067-1513) on an Agilent 2100 Bioanalyzer system.

### Quantitative RT-PCR

Reverse transcription of 500 ng of total RNA was performed using the Superscript II or IV kit and a mixture of random hexamers and oligo dT following manufacturer’s instructions at 42°C for 50 minutes followed by inactivation at 70°C for 15 minutes (Life Technologies). Primers were designed using Primer Blast and Primer3 (Supplementary Table. 1). Real-time PCR amplification was performed on a LightCycler 480 (Roche) using the SYBR green master mix. All PCR were performed using a standardized protocol and data were analyzed with the Lightcycler 480 software version 1.5.0.39 (Roche). Primer efficiency was determined by absolute quantification using a standard curve. For each sample, fold-change was obtained by comparative quantification and normalization to expression of the *HPRT, GAPDH and PPIA* (*DUX4* and 4q35 genes) *or 36B4* (characterization of hiPSCs) housekeeping genes used as standard. Data are expressed as means ± SEM.

### 3D FISH

Cells were grown on 4-well slides (Millipore) or 4-well PCA slides (hIPSCs; Sarstedt), fixed in 4% paraformaldehyde and treated as described (Ottaviani et al., 2009) to maintain the 3D structure of the cells. All probes were denatured at 80*±*1°C for 5 minutes before hybridization. Nuclei were counterstained with DAPI (Sigma) diluted to 1 mg/ml in PBS and mounted in Vectashield (Vector Laboratories). Probes were either purchased (RainbowFISH, Cytocell) or produced by nick translation using after-mentioned BAC clones (CHORI) as template: RP11-521G19 (D4S139); RP11-33M11 (*FAT1*); RP11-159A22 (*SORBS2*); RP11-374K13 (*ASCL1*) and RP11-451F20 (*WWC2*), respectively.

Images were acquired using a confocal scanning laser system from Zeiss (Germany). A 63x Plan-APOCHROMAT, oil immersion, NA 1.40 objective (Zeiss) was used to record optical sections at intervals of 0.24µm. The pinhole was set the closest to 1 Airy with optical slices in all wavelengths with identical thickness (0.4µm). Generated .lsm files with a voxel size of 0.1µm x 0.1µm x 0.24µm were processed using the IMARIS software (Bitplane, AG). After 3D analysis, we compared the mean volume ratio of nuclei and of the probe signals. Values were compared using a non-parametric Kruskal-Wallis test; population of separated or adjacent signals were compared to the control condition using a Chi-square test.

## Supplementary information

**Supplementary table 1.**
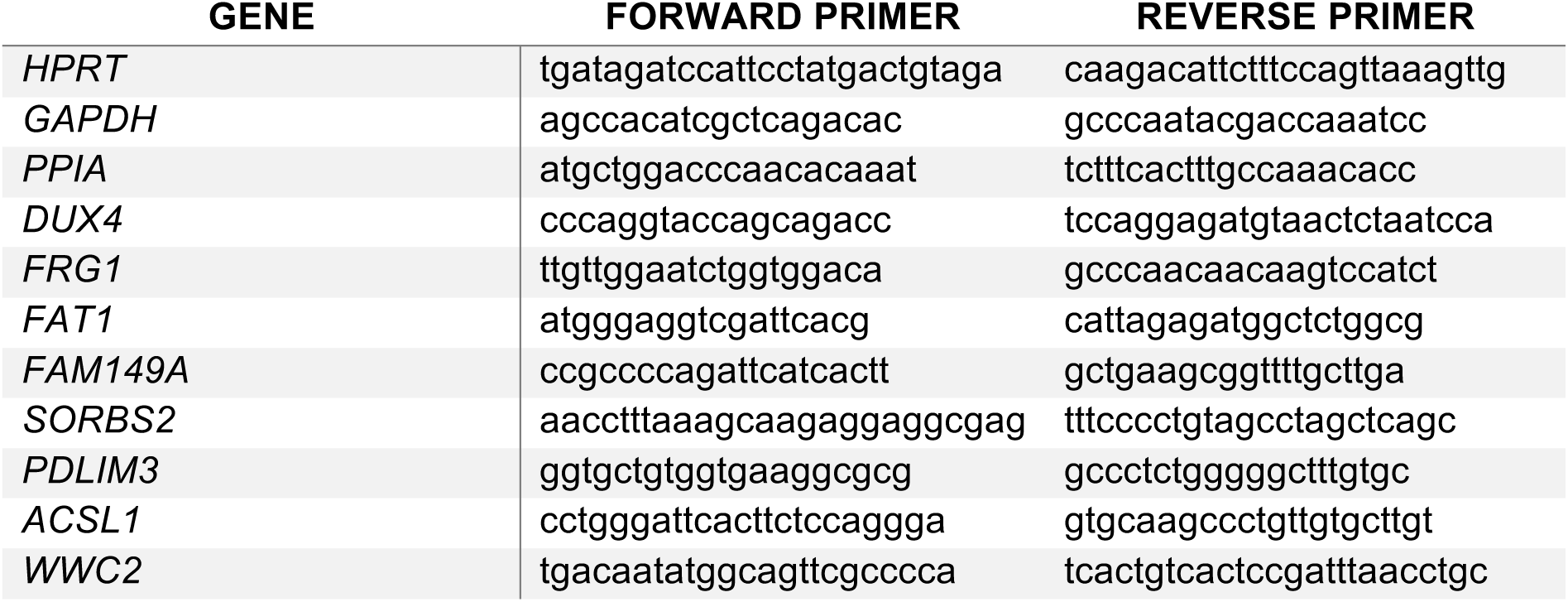
Sequence of the primers used for RT-qPCR.

**Supplementary figure 1.**
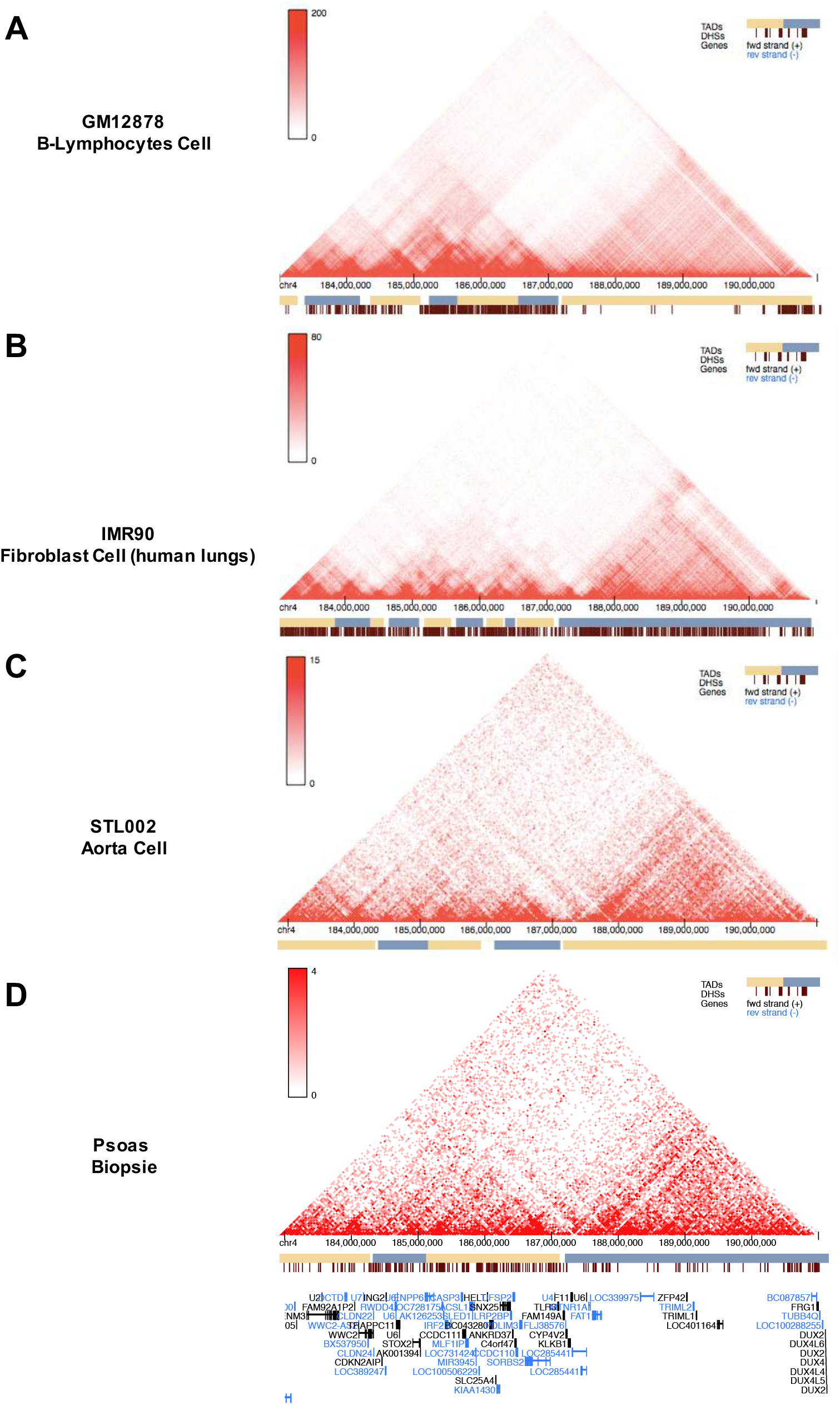
Chromatin conformation capture of the 3D organization of the 4q35 locus (last 7Mb). Representative heat maps of the last 7Mb of the 4q35 locus, revealed in other studies by HiC or ChIA-PET (STL002) and available in the “3D Genome Browser” (ENCODE project, (Y. Wang et al., 2018)). **A.** As shown by the scale intensity (200 and 80; respectively), we present the two heat maps with the highest resolution for Lymphocytes (GM12878) and fibroblasts (IMR90). **B.** Data from the most relevant tissue found in the database are also presented (e.g., muscle related cells/biopsy). The two sets of data were generated from a ChIA-PET (STL002, aorta cells) and a HiC (*Psoas* muscle biopsy) assay with a low resolution (15 and 4; respectively). Each map is presented with the corresponding Topologically Associated Domains (TADs) and DNase I hypersensitive sites (DHS) when available. The most subtelomeric TAD (stopping between the *FAT1* and *SORBS2* region) is conserved between samples.

**Supplementary figure 2.**
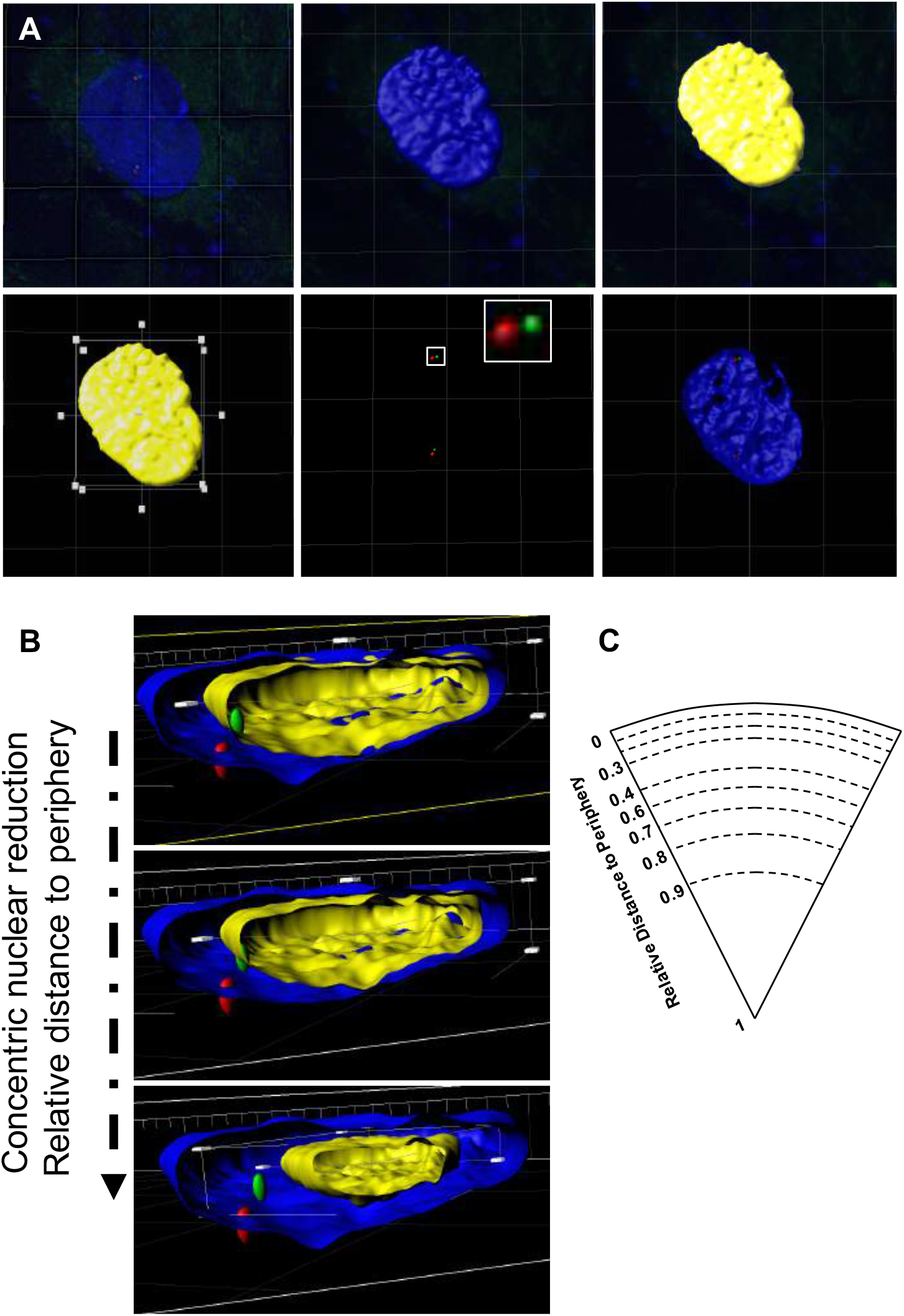
Processing of 3D reconstructed nuclei. Nuclei with regular ellipsoidal shape were selected and each nucleus was analyzed independently. **A.** Thresholded DAPI and probe signals were acquired by confocal microscopy (*z*-stacks of 0.24µm thickness). Three-dimensional objects were created for each. **B.** A reduced reproduction (yellow structure) of the object corresponding to the nuclear rim (Blue) was generated having the same center defined as the center of the cuboid in which that object is inscribed, with dimensions reduced until the FISH signal matches the reduced envelope. **C**. For each nucleus, this reduction of dimensions was mathematically turned into equivalent sphere volume reduction in order to obtain a distribution of the FISH signal within the nuclear volume for each experimental condition.

**Supplementary figure 3.**
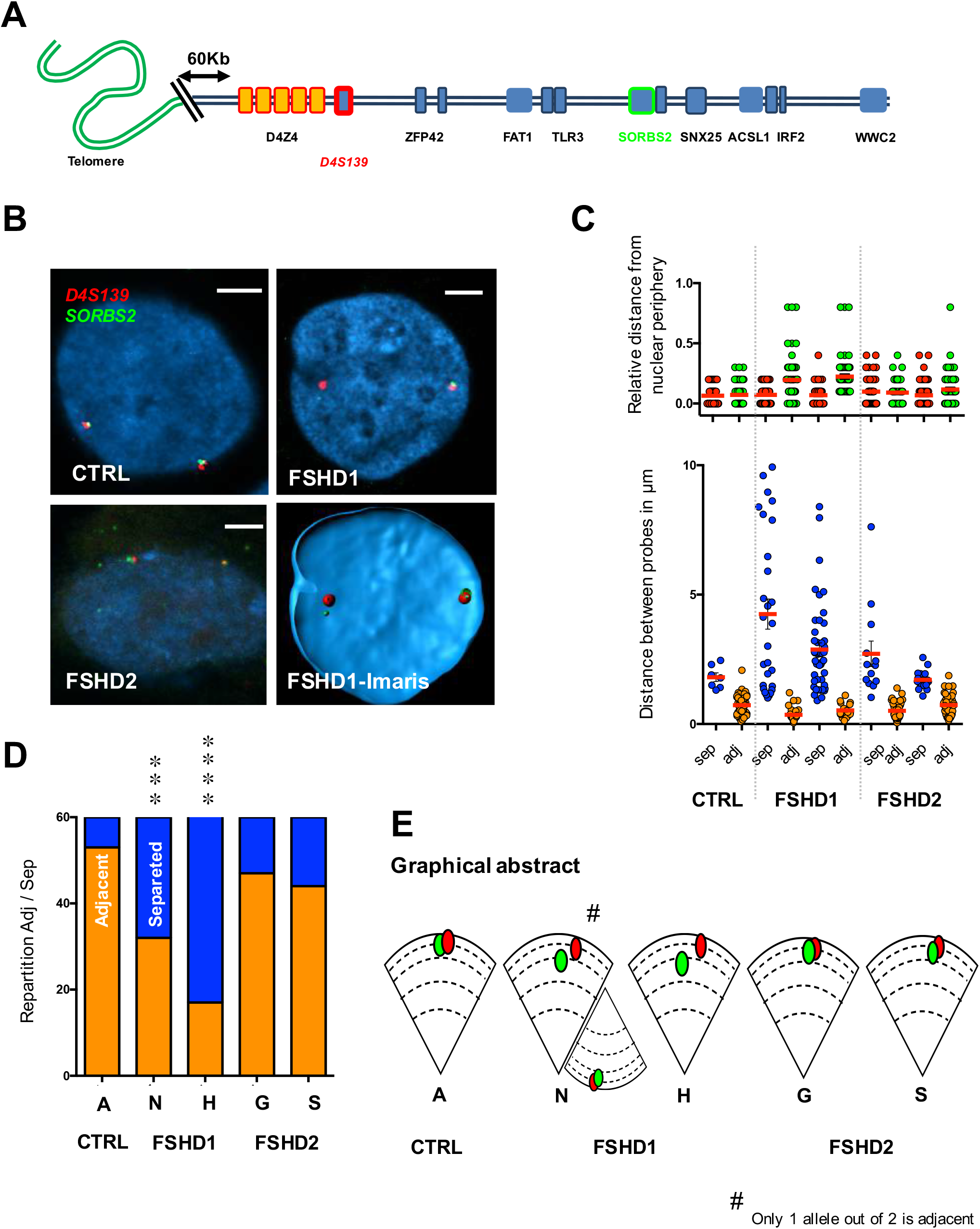
3D DNA FISH between the distal 4q35 region and the *SORBS2* gene. **A.** Graphical representation of the last 7Mb of the chromosome 4q arm, from the telomere (green) to the *WWC2* gene (blue square). 3D FISH was done using two sets of probes corresponding either the D4S139 VNTR (red) or the *SORBS2* region (green). **B.** Representative pictures and 3D reconstruction using IMARIS are presented along with their associated quantifications. Scale is indicated by a white bar (=2µm) **C.** We measured the distance of each probe to the nuclear periphery and distance between the probes. **D.** For each cell, histogram displays the percentage of separated or adjacent signals (sep, adj; respectively). Statistical significance was determined using a Chi-Square test (n=30 cells per sample, 60 alleles). **E.** Distribution of the average distances between probes and their respective distances to the periphery in each cell line. In one FSHD1 cell line (N), signals were adjacent for only one out of two probes (85% of cells), likely corresponding to the uncontracted allele. Frequencies of adjacent signals are decreased in all FSHD1 cells. *** *p* < 0.005; **** *p* < 0.001. Signals are mostly found at the periphery regardless of disease status.

**Supplementary figure 4.**
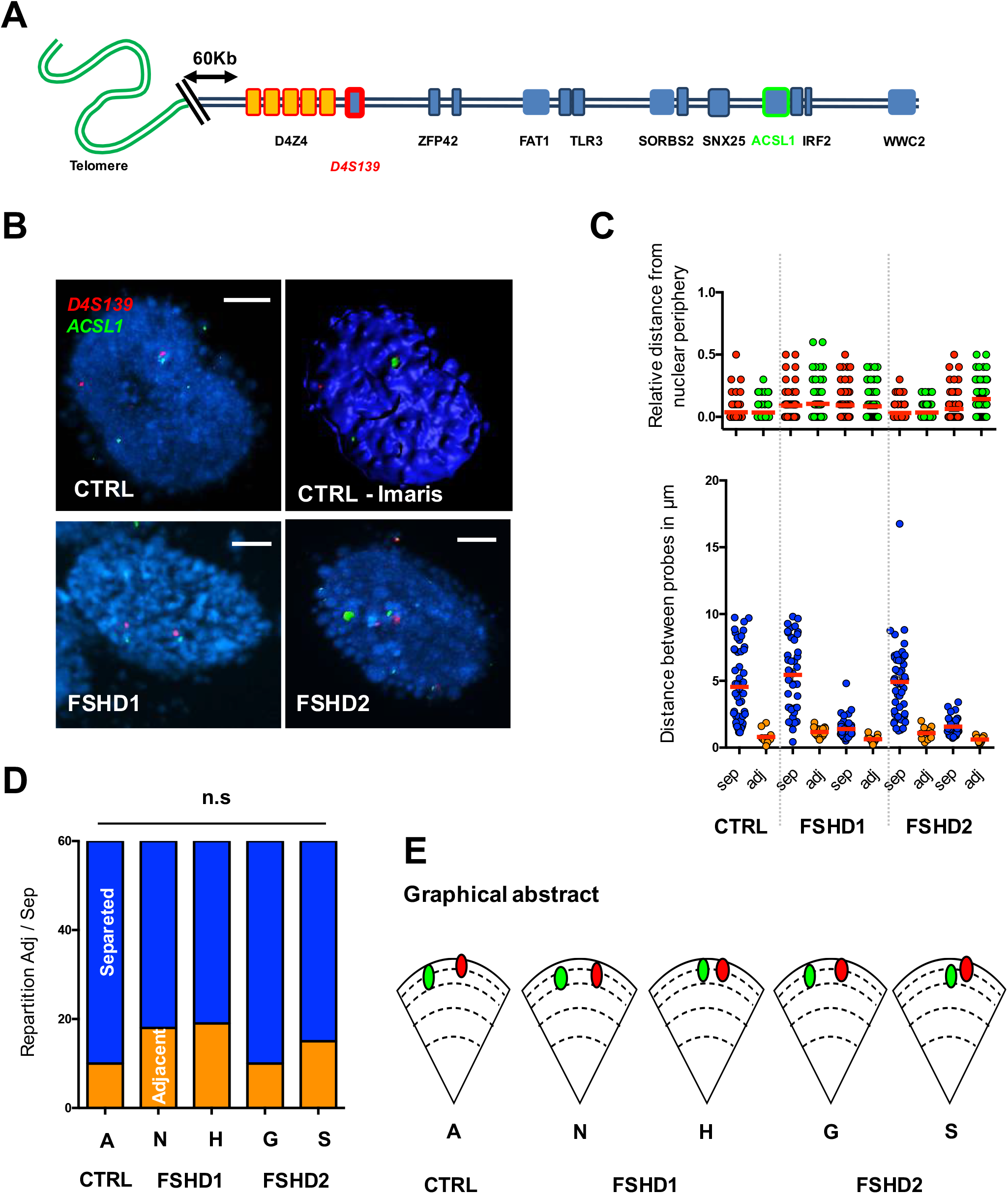
3D DNA FISH between the distal 4q35 region and the *ACSL1* gene. A. Graphical representation of the last 7Mb of the chromosome 4 long arm, from the telomere (green) to the *WWC2* gene (blue square). 3D FISH was done using two sets of probes corresponding either the D4S139 region (red) or the *ASCL1* locus (green). **B.** Representative pictures and 3D reconstruction using IMARIS along with their associated quantifications. Scale is indicated by a white bar (=2µm) **C.** We measured the distance of each probe to the nuclear periphery and distance between the probes. **D.** For each cell, we evaluated if signals were separated or adjacent (sep, adj; respectively). Statistical significance was determined using a Chi-Square test (n=30 cells per sample, 60 alleles). **E.** Graphical representation of the average distances between probes and their respective distances to the periphery in each cell line. Signals are mostly found at the periphery regardless of disease status. Localization between the different signals is not statistically different between samples.

**Supplementary figure 5.**
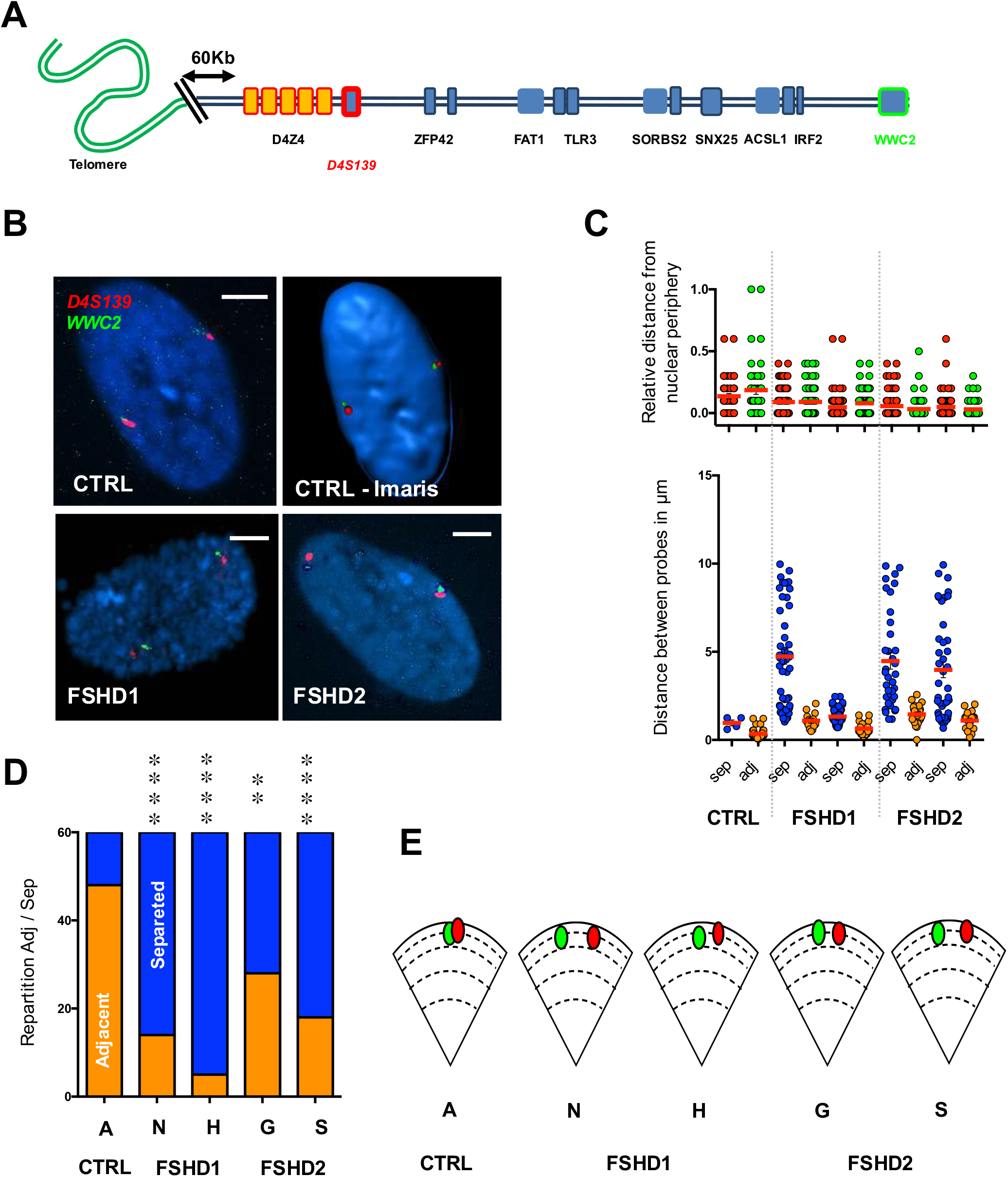
3D DNA FISH between the distal 4q35 region and the *WWC2* gene. **A**. Graphical representation of the last 7Mb of the chromosome 4 long arm, from the telomere (green) to the *WWC2* gene (blue square). 3D FISH was done using two sets of probes corresponding either the D4S139 (red) or the *WWC2* region (green). **B**. Representative pictures and 3D reconstruction using IMARIS along with their associated quantifications. Scale is indicated by a white bar (=2µm) **C**. We measured the distance of each probe to the nuclear periphery and distance between the probes. **D**. For each cell, we evaluated if signals were separated or adjacent (sep, adj; respectively). Statistical significance was determined using a Chi-Square test (n=30 cells per sample, 60 alleles). **E**. Graphical abstract of the average distances between probes and their respective distances to the periphery in each cell type. Signals are mostly found at the periphery regardless of disease status. Frequencies of adjacent signals are decreased in all FSHD cells. ** *p* < 0.01; **** *p* < 0.001.

**Supplementary figure 6.**
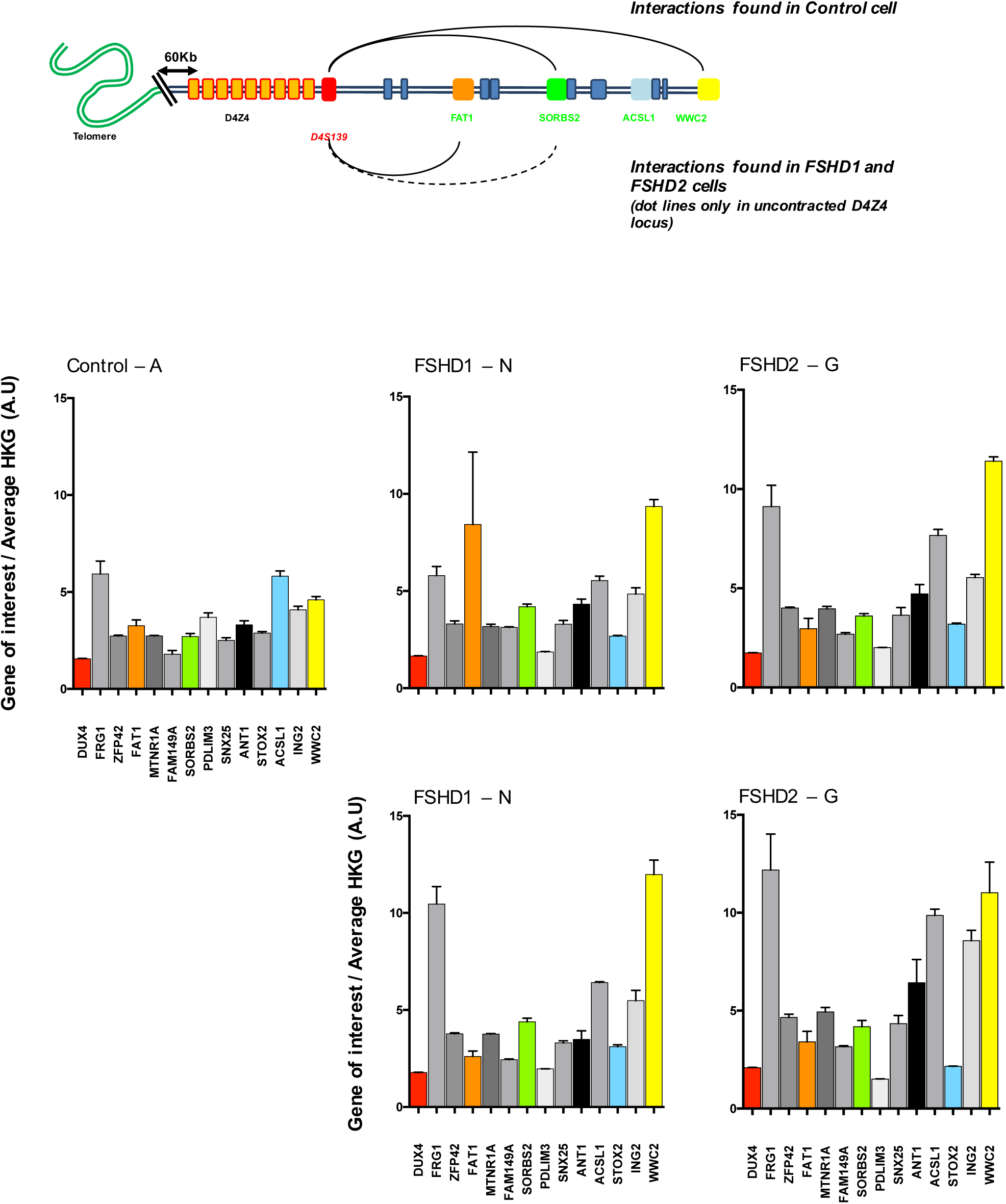
3D genome organization within the 4q35 locus and gene expression. Schematic representation of interactions described by 3D FISH experiments (top) and associated gene expression (bottom). Each gene region processed by 3D FISH is represented by a given color (e.g., *FAT1*, orange; *SORBS2*, green; *ACSL1*, blue; *WWC2*, yellow). We report the same RT-qPCR assay as in Figure. 3 but using only normalization to House Keeping Genes (HKG: *HPRT, PPIA* and *GAPDH*; Gene/HKG ratio is represented) in control (A) FSHD1 (N, H) and FSHD2 (G, S) cells. Each measure represents the average fold-change expression of six independent assays (biological triplicate in technical RT duplicate). Ratios are reported per cell line and organized regarding to their genomic localization in order to reveal potential domains (as domain of expression ruled by similar epigenetic markers and chromatin organization).

## Acknowledgements and Funding

We are indebted and thank all patients for participating in this study. We acknowledge the Marseille Stem Cell core facility for technical contribution. This study was funded by “Association Française contre les Myopathies” (AFM; MNHDecrypt and TRIM-RD), Agence Nationale de la Recherche (ANR-13-BSV1-0001). CD and JM were the recipient of a fellowship from the French Ministry of Education and FSH Society. CL is the recipient of a fellowship from the French Ministry of Education. MCG was the recipient of a fellowship from the FSH Society. CB was the recipient of fellowship from the Algerian Ministry of Education and Fondation pour la Recherche Médicale.

## Authors’ contribution

Drs. Marie-Cécile Gaillard, Natacha Broucqsault, Camille Dion, Cherif Badja, Julia Morere, Stéphane Roche and miss Camille Laberthonnière set up the experimental conditions, performed the experiments analyzed and organized the data. These authors report no disclosure.

Dr. Karine Nguyen provided and processed patients’ samples. Dr. Nguyen reports no disclosure.

Dr Jérôme D. Robin: Contributed to the design and coordination of the project, analyzed the data, performed the statistical analysis and wrote the paper. Dr. Robin reports no disclosure. Dr. Frederique Magdinier: designed and coordinated the project, provided funding, performed the statistical analysis, analyzed the data and wrote the paper. Dr. Magdinier reports no disclosure.

## Graphical Abstract

As found in the literature the 4q35 locus is composed of several Lamina associated domains (LADs) and Topological active domain (TAD). The locus is also linked to Facio Scapulo Humeral Dystrophy (FSHD). Modification of this chromatin organization in FSHD cells remains elusive. Using 3D DNA FISH probes (color-coded gene box) across the locus at strategic places, we established a first 3D organization across the 4q35 locus. Further, we found that the looser conformation found in FSHD cells is associated with increased gene expression (bottom panel, increased expression represented with bolder font; Grey: overexpressed in FSHD cells; Black: over expressed in FHSD1; red: only over expressed in FHSD2). Using hIPSC we found that the 3D organization is set to a new chromatin organization regardless of the disease status.

Altogether, our data reinforce current hypothesis for the etiology of FSHD, linked to defect in differentiation and bring new results for the variegated expression associated with the variability in disease penetrance between patients

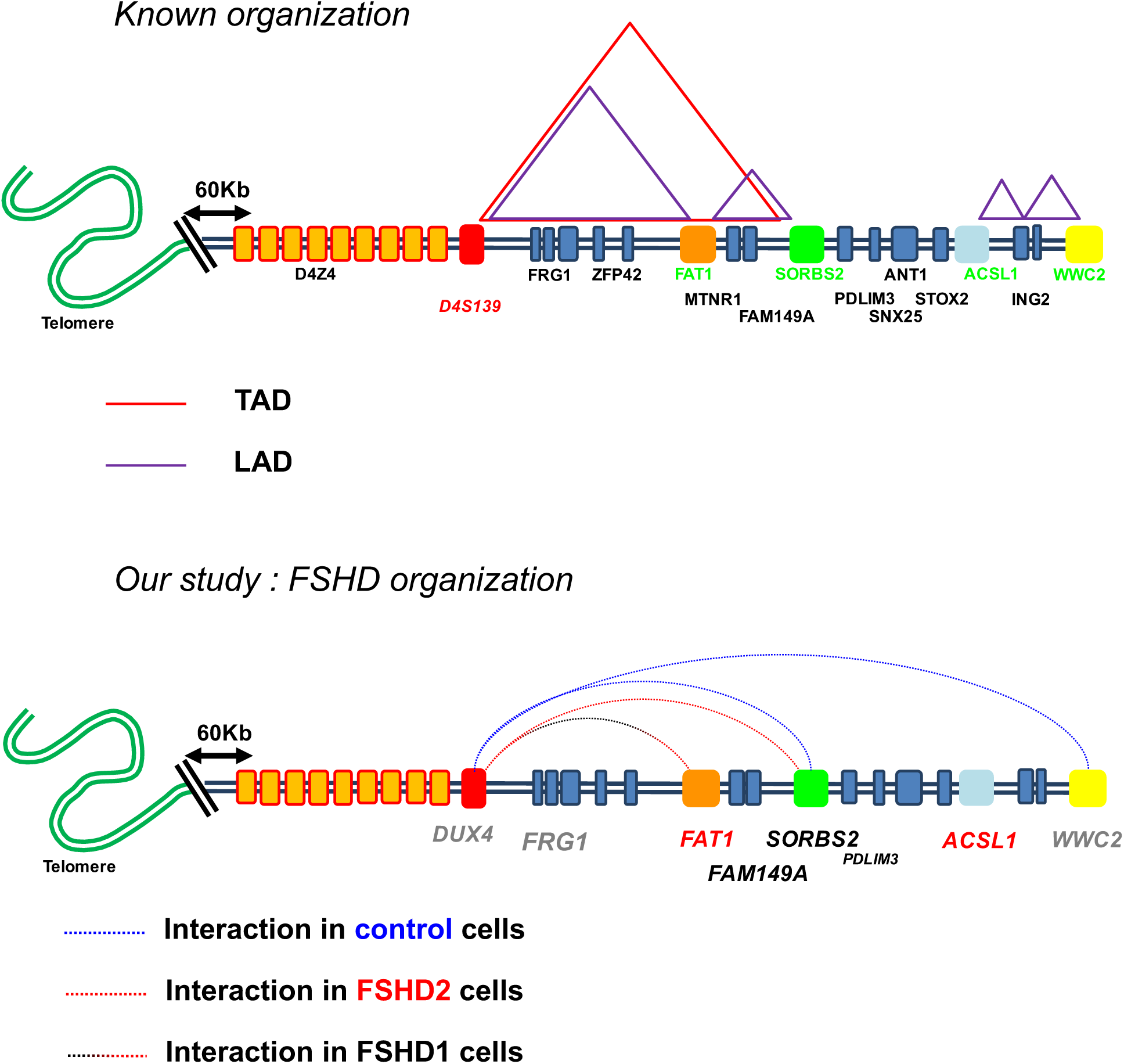

